# Prevalence and zoonotic risk of multidrug-resistant Escherichia coli in bovine subclinical mastitis milk: Insights into the virulence and antimicrobial resistance

**DOI:** 10.1101/2024.08.22.609233

**Authors:** Tonmoy Chowdhury, Mithu Chandra Roy, Ferdaus Mohd Altaf Hossain

**Affiliations:** Department of Dairy Science, Faculty of Veterinary, Animal and Biomedical Sciences, Sylhet Agricultural University, Bangladesh; Department of Microbial Biotechnology, Faculty of Biotechnology & Genetic Engineering, Sylhet Agricultural University, Bangladesh

**Keywords:** Escherichia coli, Shiga Toxin, Antimicrobial Resistance, Subclinical Mastitis, Zoonosis

## Abstract

The emergence of antibiotic-resistant microorganisms has made AMR a major issue on a global scale, and milk has the potential to be a possible source for the propagation of resistant bacteria causing zoonotic diseases. SCM cases, often overlooked and mixed with normal milk in dairy farms, frequently involve *E. coli*, which can spread through contaminated milk. We conducted this study to determine the prevalence of virulence genes, ARGs, antimicrobial susceptibility, and the genetic relatedness of MDR-STEC isolated from SCM milk. SCM-positive bovine milk was subjected to *E. coli* detection using cultural, biochemical, and molecular methods. Further, we detected STEC virulence genes including *stx1, stx2,* and *eaeA*. STEC isolates were tested for ARGs including *blaSHV*, *CITM*, *tetA,* and *aac(3)-IV*, and underwent AST. Moreover, we performed a phylogenetic analysis of the *stx1* gene of MDR-STEC. SCM was detected in 47.2% of milk samples of which 50.54% were *E. coli* positive. About 17.20% of *E. coli* isolates contained STEC virulence genes, and *stx2* was the most prevalent. Moreover, all STEC isolates harbored at least one of the ARGs, while about 43.75% of the isolates carried multiple ARGs. Additionally, all the STEC isolates showed multidrug resistance, and were found to be fully resistant against amoxicillin, followed by ampicillin (87.50%) and gentamycin (75%); and were mostly sensitive to aztreonam (81.25%) and meropenem (68.75%). We found that MDR-STEC isolated from SCM were closely related to disease-causing non-O157 and O157 strains of different sources including cattle, humans, and food. These findings indicate a significant zoonotic risk.

## Introduction

Antimicrobial resistance (AMR) has currently become a matter of worldwide concern in recent decades because of the rapid emergence of antimicrobial-resistant bacterial strains, particularly *Escherichia coli* (*E. coli*), a rod-shaped gram-negative bacteria, which is considered a threshold threat to humans, animals, and the environment worldwide, including Bangladesh (Al Amin et al., 2020; Hossain et al., 2011; Rousham et al., 2018). The Shiga toxin-producing *E. coli* (STEC) serovars which contain important cytotoxins such as Shiga toxin-1 (*stx1*), Shiga toxin-2 (*stx2*), Intimin (*eaeA*), and many more, address intense concerns regarding food safety, antibiotics treatment failure in STEC-causing diseases, and finally ensuing AMR causing animal and public health hazard (Ansharieta et al., 2021; Farrokh et al., 2013). *E. coli* is considered one of the most important opportunistic bacteria that has been associated with antibiotic resistance and sub-clinical mastitis (SCM) (Hinthong et al., 2017). Additionally, the potential risk of *E. coli* in SCM, and the presence of harmful STEC have been reported (Ahmadi et al., 2020; El-Khabaz et al., 2022). Furthermore, excessive antibiotic usage in dairy cows has a substantial impact on the emergence of multidrug resistance in STEC isolates available in milk and dairy products, which soon could result in serious AMR scenarios (Ahmadi et al., 2020; Daini et al., 2005; Mylius et al., 2018).

Cattle itself and raw milk are considered to be reservoirs for STEC bacteria and accountable for STEC *E. coli* O157:H7 infection in humans (Ansharieta et al., 2021; Elafify et al., 2020). Moreover, other disease-causing non-O157 serotypes of STEC can also be found in bovine milk (Elhadidy & Mohammed, 2013; Farrokh et al., 2013). The presence of STEC has also been reported in bovine clinical mastitis (visible signs and symptoms) and SCM (without visible signs and symptoms) milk as well (Ahmadi et al., 2020; El-Khabaz et al., 2022; Momtaz et al., 2012). Usually, hemolytic uremic syndrome (HUS) and hemorrhagic colitis (bloody diarrhea) are the two most common illnesses that occur due to STEC. About 90% percent of STEC-infected patients develop hemorrhagic colitis, while 5% to 15% suffer HUS (Gyles, 2007; Smith et al., 2014; Tarr et al., 2005). STEC-causing HUS outbreaks in children associated with milk and milk product consumption were reported in France and England (Jones et al., 2019; Treacy et al., 2019). In Bangladesh, enterotoxigenic *E. coli* (ETEC), which is closely linked to STEC (Kaper et al., 2004; Nataro & Kaper, 1998), was estimated to be responsible for 10% – 20% of pediatric diarrhea (Qadri et al., 2005). Hence, the presence of STEC in consumable milk and milk products can seriously harm human health.

Moreover, in mastitis, *E. coli* is considered the most common environmental bacterial infection (Eberhart, 1984; Goulart & Mellata, 2022), and also STEC has been reported to be an emerging causal pathogen of bovine mastitis (Murinda et al., 2019). Additionally, the prevalence of multidrug-resistant (MDR) STEC tends to rise over time (Smith et al., 2014). STEC isolated from bovine mastitis milk already shows resistance against penicillin, streptomycin, tetracycline, ampicillin, gentamycin, etc. antibiotics (Ahmadi et al., 2020; Momtaz et al., 2012). STEC isolated from bovine mastitis milk had been found harboring antibiotic resistance genes (ARGs) such as beta-lactam class resistance genes (*blaSHV, CITM*, etc.), tetracycline resistance genes (*tetA* and *tetB*), aminoglycoside resistance gene (*aac(3)-IV*), and many more (Momtaz et al., 2012). These reports raise concerns about not only the safety of consumable milk and milk products but also bovine udder health which is essential for successful dairying and safe milk production.

Furthermore, SCM in dairy cattle is predominant in Bangladesh and mostly goes unnoticed by the farmer, as a result, the SCM milk gets mixed with normal raw milk in commercial dairy which then goes to market for human consumption (Das et al., 2020; Md. S. Hasan et al., 2021; Kahir et al., 2008; M. M. Rahman et al., 2010). This increases the chance of SCM milk containing STEC getting into the human food chain. In Bangladesh, no study has been conducted on STEC in bovine SCM milk, and minimal data are available on MDR-STEC and their ARGs concerning bovine SCM milk. So, there is still a huge gap in knowledge on MDR-STEC in bovine SCM milk. In addition, there is a lack of organized and standard regulations to govern safe milk supply and production. Hence, we intended to monitor the MDR-STEC scenario for the bovine SCM milk. These considerations led us to conduct this study to detect the prevalence of STEC in bovine SCM milk. We also identified their virulence and antimicrobial resistance genes, analyzed their antimicrobial resistance profiles, and performed a phylogenetic analysis based on the *stx1* gene to determine their genetic relatedness to other strains.

## Materials and methods

### Sample collection

The samples were collected from 13 upazilas of Sylhet district, Bangladesh (**Figure 1**). A total of 39 dairy farms from 13 different upazilas (3 farms in each upazila) were randomly selected for the collection of raw milk samples. A total of 390 raw milk samples from individual cows (crossbreed) of randomly selected dairy farms were collected (**Figure 1**). Milk from all these cows was actively being sold to the market for consumption and dairy products manufacturing purposes. Before collecting the raw milk samples, the milking man’s hand and udder were properly cleaned with soap and clean water. The raw milk from all four quarters was then taken directly into a sterilized 15 ml falcon tube. A small amount of raw milk was taken out with a sterilized plastic dropper for conducting the SCM test (Whiteside test). Alongside, the falcon tubes were stored in an airtight icebox and transported as soon as possible to the Dairy Science Laboratory, Department of Dairy Science, Sylhet Agricultural University, Bangladesh for further experiments.

**Figure 1:**
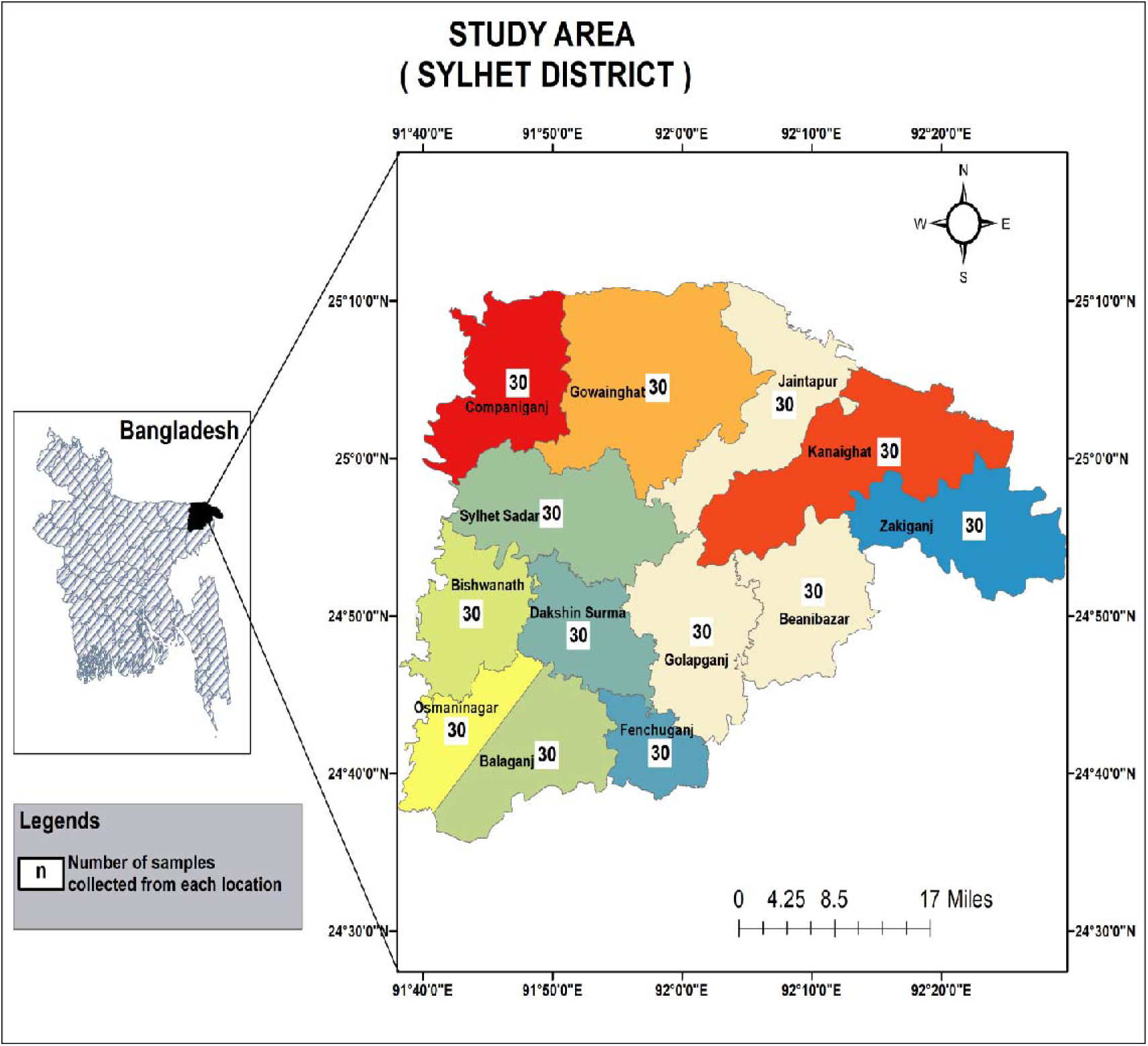
Map of the study area. (A total of 30 milk samples were collected from each upazila. A total number of 390 milk samples were collected from Sylhet district, Bangladesh)

### Subclinical mastitis (SCM) detection

To detect SCM, the Whiteside test (WST) was performed immediately after the collection of the milk samples at farms. The Whiteside test (WST) was performed following the descriptions of previous studies (Astermark et al., 1969; Kahir et al., 2008). About 5 drops of milk sample from a falcon tube were taken on a glass slide with a sterilized plastic dropper. Two drops of normal sodium hydroxide (NaOH) solution (4%) were then added to the milk sample on the glass slide. The solution was gently mixed using a glass stirrer. After 20-25 seconds, if there was the presence of gel formation and/or the solution was broken down into small white flakes, the sample was considered as a sub-clinical mastitis positive.

### Bacterial culture

Pre-enrichment of *E. coli* was done at 37°C for 24 hours in Tryptone Soya Broth (TSB) (HiMedia, India). TSB acts as a good medium for enrichment for *E. coli* and has been used in previous studies as well (Reinders et al., 2002; Wallace et al., 1997). For the isolation of the *E. coli* colony, we used Eosin Methylene Blue (EMB) agar (Oxoid Ltd, United Kingdom).

EMB agar has been used widely because of its selectivity for gram-negative bacteria and differential characteristics for *E. coli* where the medium develops colonies with green metallic sheen in the presence of *E. coli* (Antony et al., 2016; Fahim et al., 2019). The inoculated EMB agar was incubated for 24 hours at 37°C. Bacterial colonies showing a dark center with a green metallic sheen were presumptively considered *E. coli* colonies. Repeated culture of these presumptive *E. coli* colonies was performed in EMB agar following previously described incubation time and temperature until we got a pure single colony. These single colonies of *E. coli* were further subjected to gram staining and biochemical tests.

### Gram Staining of suspected E. coli colony

A single drop of sterilized distilled water (DW) was taken on the glass microscope slide. Then one single suspected *E. coli* colony was taken with the help of a sterilized platinum loop and mixed with the water drop on the glass microscope slide to dilute the bacterial concentration. After that, the sample was heat-fixed by passing the glass slide 3-4 times over the flame of the spirit lamp. Next, the gram staining was done following the procedures of TJ Beveridge (2001) (Beveridge, 2001). Finally, the slide was observed at 100X under a light microscope (Model: G-260, Optima, Taiwan).

### Biochemical tests for E. coli

We performed the Motility Indole Urea test (MIU), Methyl Red and Voges-Proskauer test (MR-VP), Citrate utilization test (CU), Sugar fermentation test, according to the procedures described by (Feng et al., 2020), and catalase test according to (Karas et al., 2007). The positive result for indole and motility and a negative result for urease was considered as an *E. coli* positive sample (Purkayastha et al., 1970). Development of pink or red color in the medium on the Methyl-Red (MR) test was considered as *E. coli* positive characteristics (Karim et al., 2020) while *E. coli* giving a negative reaction (no color development) for the VP test was considered as *E. coli* positive (Karim et al., 2020). As the development of blue color in the CU test indicates a positive growth reaction, so *E. coli* showing a negative reaction (no color development) in CU was considered as an *E. coli* positive sample (Dash et al., 2012). Moreover, the production of gas and media color changing into yellow in the Sugar fermentation test were considered characteristics of the *E. coli* positive sample (Dash et al., 2012). Furthermore, in the case of the catalase test, if there was a presence of bubbling within the 1-minute observation time then the sample was considered catalase positive, and if there was no bubbling throughout the one-minute observation, the sample was interpreted as catalase negative.

### Preservation of E. coli isolates

The EMB culture samples that showed characteristics of *E. coli* and confirmed in gram staining and biochemical tests were preserved for further use. Preservation was done using nutrient broth (NB) and glycerol (15%) mixed solution described by a previous study (Gorman & Adley, 2004). To stock the bacterial cultures, firstly Nutrient broth (HiMedia, India) was prepared by dissolving at an amount of 13 gm in 1000 ml DW and autoclaved at 121°C for 15 minutes at 15 psi of pressure for sterilization. After completing the autoclave process, the nutrient broth was kept in a water bath to cool down at a temperature of 37°C and distributed at an amount of 10 ml in each test tube. Then a single colony of *E. coli* from EMB agar was streaked into sterilized nutrient broth containing test tubes. The test tubes were closed using cotton plugs and then incubated at 37°C for 24 hours. After incubation, the growth of bacteria was confirmed by the development of turbidity in the broth. Finally, for preservation, 850 µl of broth culture with 150 µl of sterile 100% glycerol was kept in a sterilized 1.5 ml Eppendorf tube and gently inverted 2 times to mix. The Eppendorf tubes were marked with respective sample IDs and kept at −20°C for future use.

### Bacterial DNA extraction

Extraction of bacterial DNA was done by the boiling method as described in a previous study (Md. S. Islam et al., 2021). However, we slightly modified some steps in the method but didn’t alter the basic steps of boiling method DNA extraction (Md. S. Islam et al., 2021). Firstly, Nutrient agar (NA) (HiMedia, India) was prepared by adding and dissolving NA at an amount of 28 gm in 1000 ml DW in a sterilized conical flask. The medium was then autoclaved at 121°C for 15 minutes at 15 psi of pressure. After autoclaving, the conical flask was kept in a water bath to cool down to 45°C and then spread on a petri dish inside a laminar airflow cabinet (Biobase, China). When the NA medium solidified in the petri dish, then the preserved cultures of *E. coli* isolates were streaked on the NA medium using a sterilized platinum loop. The petri-dish were then incubated at 37°C for 24 hours. After the incubation period, the medium-sized colonies were taken into nuclease-free water (500 µl) in an Eppendorf tube (Hyclone Laboratories Inc, USA) and the Eppendorf tube was vortexed for 10 seconds using a vortex machine (Model: MS-S, DLAB, China). After that, to pellet the contents of Eppendorf tubes, centrifugation was done at 13000 rpm for 5 minutes using a centrifuged machine (Model: D3024, DLAB, China). The supernatants were discarded from the Eppendorf tubes. The tubes were again filled with 500 µl nuclease free water and vortexed for 5 seconds. Following that, the tubes were then placed on boiling water in the water bath for 10 minutes. After that, all the Eppendorf tubes were immediately kept on - 20°C for 10 minutes. After cooling, the Eppendorf tubes were subjected to centrifugation at 13000 rpm for 5 minutes. Finally, 50 µl of supernatant from Eppendorf tube was moved to another sterilized Eppendorf tube and used as DNA template. The DNA templates were stored at −20°C for further use.

### Molecular detection of E. coli, STEC, and ARGs by PCR

PCR was conducted for molecular detection of *E. coli (16s rRNA),* STEC genes (*stx1*, *stx2*, *eaeA*), and ARGs (*blaSHV, CITM, tetA, aac(3)-IV*) respectively. PCR was performed on a thermal cycler (Model: T100, Bio-Rad Laboratories, Inc.). For the PCR reaction mixture, we used Emerald Amp MAX PCR Master Mix (2X) (Takara Bio, USA). All the primer sets were manufactured by Macrogen, Inc. (Seoul, South Korea). The primers used in our experiments were taken from previous studies (Dipineto et al., 2006; Guillaume et al., 2000; Hessain et al., 2015; Khan et al., 2002; Nourbakhsh et al., 2017; Parvin et al., 2020), and the details are mentioned in **(Table 1)**. PCR tubes (0.2 ml volume) were used for performing PCR. For preparing PCR reaction mixtures, at first PCR tubes were autoclaved and then kept inside a laminar airflow cabinet maintaining aseptic condition. The PCR reaction mixture was then prepared at a volume of 25 µl. We added 12.5 µl of master mix, 1µl (10 picomoles per 1µl concentration) of forward primer, 1µl (10 picomoles per 1µl concentration) of reverse primer, 4µl of DNA template, and 6.5µl of nuclease-free water (Hyclone Laboratories Inc, USA). The PCR conditions were set to initial denaturation at 95°C for 5 minutes and final extension at 72 °C for 7 minutes for all the primers. The number of cycles, temperature, and time for denaturation, annealing, and extension steps were different for each primer and have been stated in (**Table 1**).

**Table 1:**
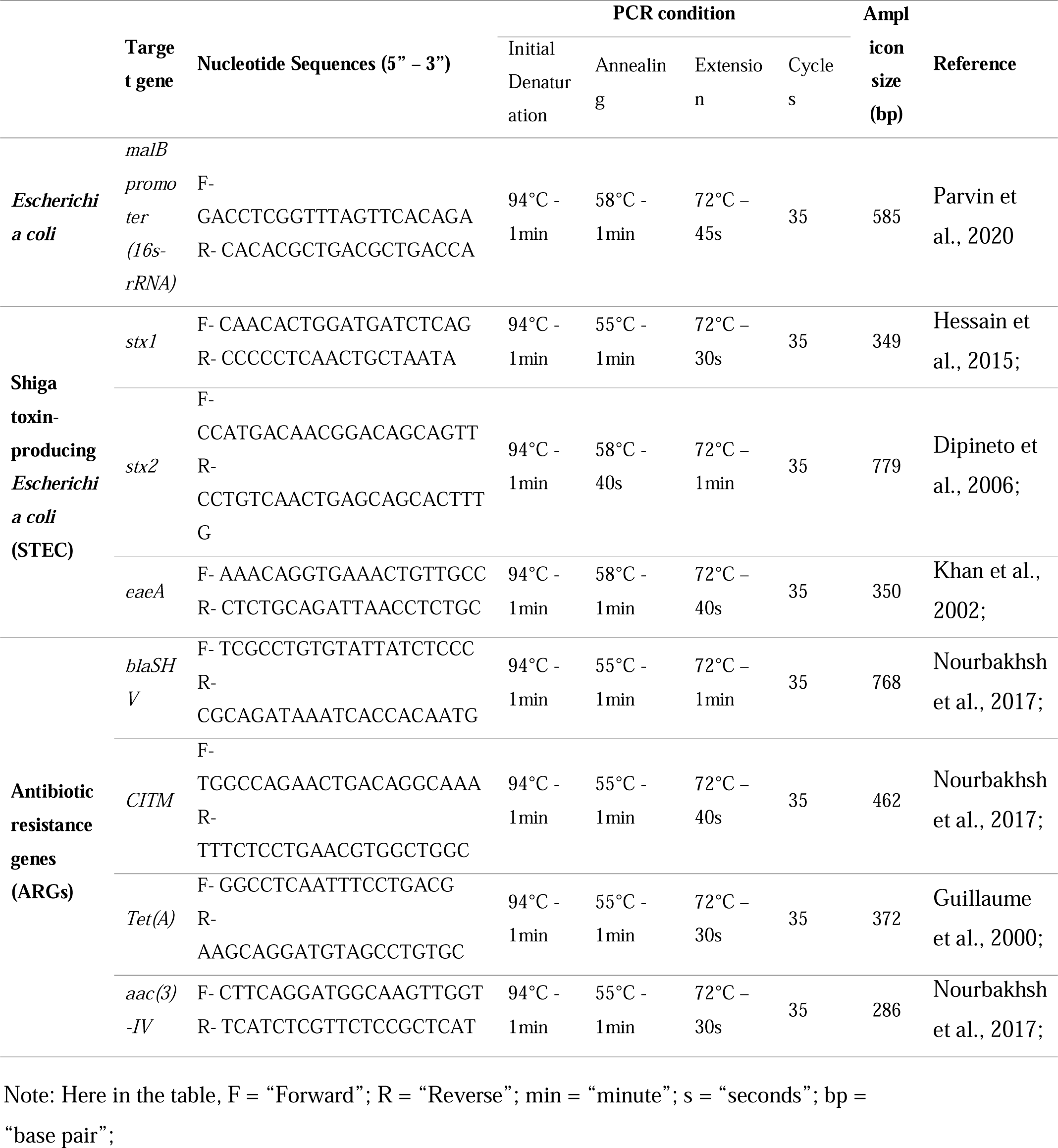
Primers used in molecular detection of E. coli, STEC, and ARGs.

### Gel electrophoresis

To perform gel electrophoresis, 50 ml of 1.5% agarose solution was prepared by mixing and dissolving 0.75 gm ultrapure agarose powder (Invitrogen, Life Technologies, USA) and 1X TAE buffer (Himedia, India). The solution was then cooled down at 60-70C and 0.5 µl of Ethidium bromide 1% solution (Carl Roth GmbH + Co. KG, Germany) was added to the agarose solution. The solution was then gently mixed and poured on the side of the gel plate where the comb was placed and left for 15 minutes to solidify. After solidification of the gel, the gel was carefully placed on the electrophoresis tank (BTLab systems, A Geno Technologies, Inc., USA), and submerged about 5-6 mm in 1X TAE buffer. The PCR products (5µl) were then loaded in the well of the gel. 5µl of 100bp DNA ladder showing each band increment from 100bp to 1000bp and a 1500bp band (Eco in action, GeneDirex, Inc., South Korea) was also loaded on a gel well. The gel electrophoresis was then performed at 100 volts for 30 minutes using a gel electrophoresis system (BTLab Systems, A Geno Technologies, Inc., USA). After completion of electrophoresis, the gel was then taken out of the gel tank and visualized under the gel documentation system using UV trans-illumination (Bio-Rad Laboratories Inc., USA).

### Antimicrobial susceptibility test

The antimicrobial susceptibility test of *E. coli* was performed following the disc diffusion method as described in previous studies and guidelines (Bauer et al., 1966; CLSI, 2020). Mueller-Hinton agar (Himedia, India) was prepared by dissolving the agar medium at an amount of 38 gm in 1000 ml of distilled water and sterilized by autoclaving at 121°C for 15 minutes at 15 psi of pressure. After autoclaving, the media was then kept in a water bath to cool down to 39°C. Then the media was poured onto a petri dish and left in the laminar airflow cabinet until the media solidified. Then colonies of *E. coli* isolates which were cultured and incubated (24 hours at 37°C) in nutrient agar (HiMedia, India), were suspended in 3ml of normal saline (0.85% NaCl solution) and matched to 0.5 McFarland standards. The samples were then dispersed using an L-shaped spreader onto the Mueller-Hinton agar medium. After that antibiotic discs (HiMedia, India) were placed on the agar medium. The Petri dishes were then kept on incubation for 24 hours at 37°C. After the completion of the incubation period, the inhibition zones were measured according to the standard guidelines of a previous study (CLSI, 2020). Bacteria were considered MDR when the bacteria showed resistance against at least 1 antimicrobial of 3 or more different classes of antimicrobials (Magiorakos et al., 2012; Rozwadowski & Gawel, 2022; Sweeney et al., 2018). We used 13 antibiotics for the antimicrobial susceptibility test **(Supplementary Table 1)**.

### PCR amplicon sequencing and phylogenetic analysis

The amplified PCR product (25 µl) of the *stx1* gene of an MDR-STEC isolate was subjected to Sanger DNA sequencing (Sanger et al., 1977). To perform phylogenetic analysis, a total of 57 nucleotide sequences of the *stx1* gene of *E. coli* isolates were retrieved from the NCBI (National Center for Biotechnology Information) database. Isolation sources of these isolates were cattle, beef, cheese, cattle feces, human, and human stool. Multiple sequence alignment was done using MEGA 11 (Molecular Evolutionary Genetics Analysis) software. The best model fit was K2P (unequal transition/transversion rates and equal base frequency), which was determined by using ModelFinder in IQ-Tree (Kalyaanamoorthy et al., 2017). The phylogenetic tree was constructed based on the maximum likelihood method on IQ-Tree version 2.3.4 (Nguyen et al., 2015). Furthermore, Fig Tree software and Microsoft PowerPoint 2019 were used for annotating the phylogenetic tree.

### Statistical analysis and data visualization

All quantitative data was recorded on Microsoft Excel 2019. The GraphPad Prism 9.3.1 statistical software was used for descriptive analysis and generating graphs.

## Results

### Prevalence of bovine SCM

The overall prevalence of bovine SCM was found about 47.2% (184 out of 390 milk samples) in Sylhet district with a lowest mean percentage of 30% in Balaganj upazila and a highest of 63.3% in Kanaighat upazila **(Figure 2a)**.

**Figure 2:**
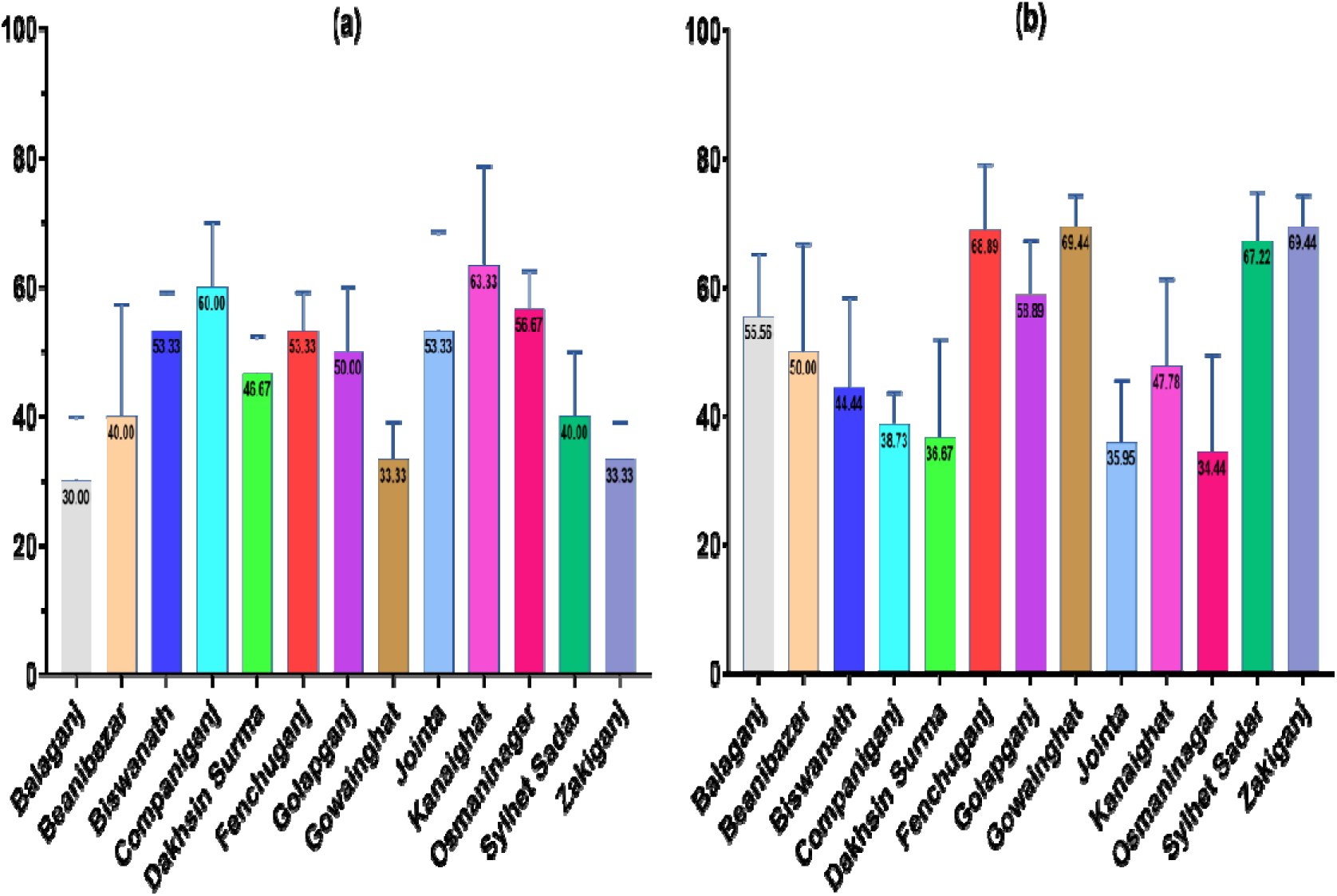
Prevalence of SCM in milk samples and detection of *E. coli* in SCM-positive samples. (The bar graph (a) shows the SCM prevalence in dairy cattle in different upazilas of Sylhet. The total number of SCM-positive samples was 184 out of 390 milk samples. The graph (b) depicts the prevalence of *E. coli* in SCM-positive samples. About 93 out of 184 SCM-positive samples were identified as *E. coli*-positive samples.)

### Prevalence of E. coli

All culture and biochemical characteristics that were found in this study for *E. coli*-positive samples are given in the supplementary file **(Supplementary Table 2).** The percentage of culture-positive presumed *E. coli* was found in 110 samples out of 184 SCM-positive samples. About 101 biochemical test-positive *E. coli* samples were found in the culture-positive samples. Among those, a total of 93 samples were found *E. coli* positive in molecular detection. Hence the total percentage of *E. coli* positive samples was 50.54% (93 out of 184) in SCM positive samples. The lowest mean percentage of *E. coli* presence in SCM milk samples was 34.44% in Osmaninagar upazila, while the Gowainghat and Zakiganj upazilas had the highest percentage of 69.44% **(Figure 2b)**.

### Prevalence of STEC and virulence genes

We found a total of 16 samples as STEC-positive. Hence, the prevalence of STEC in *E. coli*-positive samples was 17.20% (16 out of 93), and the prevalence of STEC in the SCM milk samples was 8.69% (16 out of 184 SCM samples) **(Figure 3a)**. About 31.25% (n = 5) STEC was found to contain *stx1* gene, and 56.25% (n = 9) STEC had *stx2* gene. But no sample was found positive for only the *eaeA* gene. We found 6.25% (n = 1) STEC to be harboring both *stx1* and *stx2* genes, and 6.25% (n = 1) STEC harboring both *stx2* and *eaeA* genes **(Figure 3b)**.

**Figure 3:**
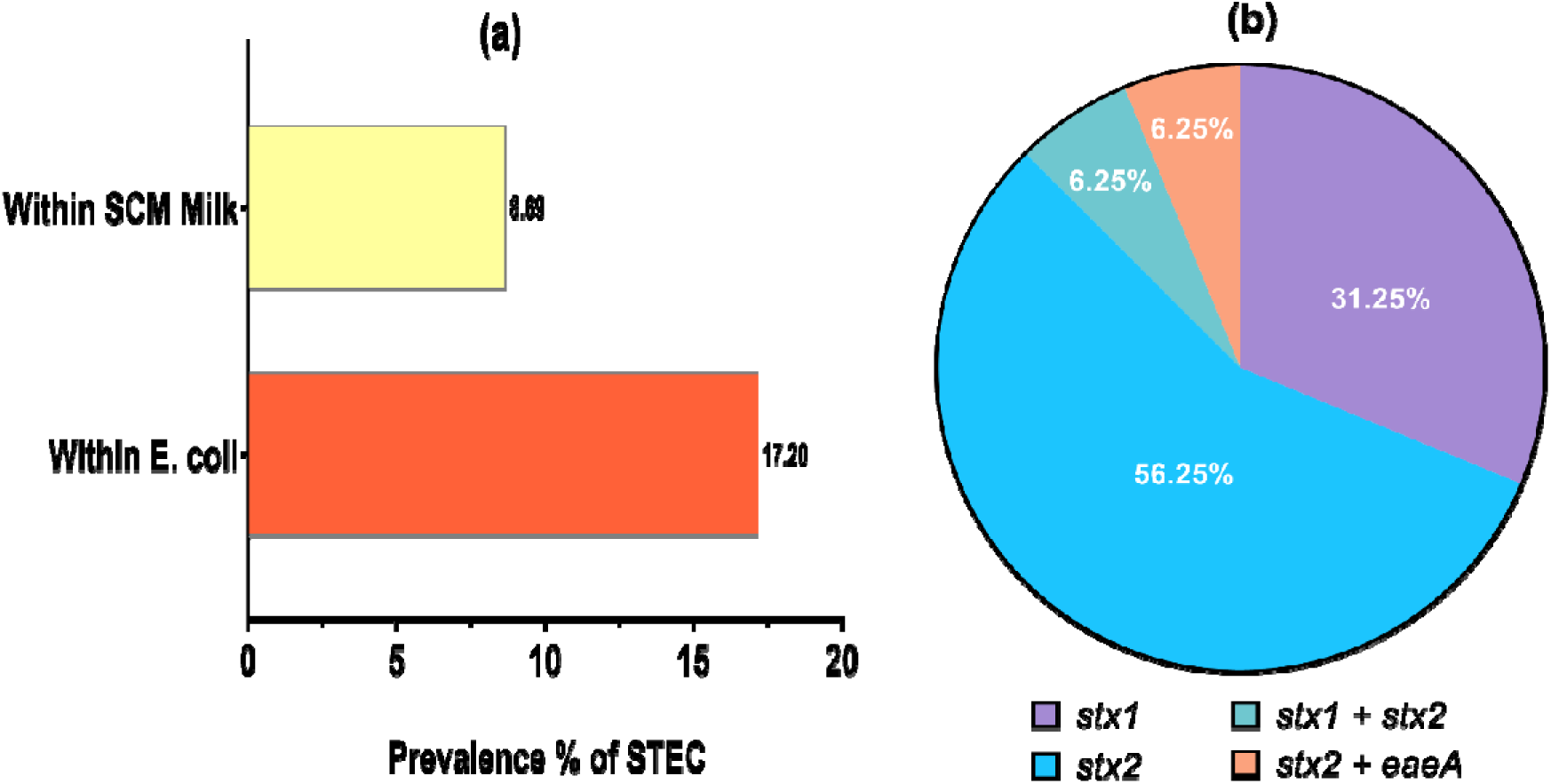
Prevalence of STEC and detection of virulence genes in STEC. (Graph (a) illustrates the prevalence of STEC in *E. coli*-positive samples (n = 93) and SCM milk samples (n = 184). The pie chart (b) shows the percentages of virulence genes found in STEC samples (n = 16).)

### Molecular detection of ARGs in STEC

In case of ARGs, all STEC isolates (n = 16 out of 16) carried at least one of the ARGs, and among them 25.00% (n = 4) harbored *blaSHV* gene, 37.50% (n = 6) contained *CITM* gene, 43.75% (n = 7) had *tetA* gene, and 50.00% (n = 8) carried *aac(3)-IV* gene **(Figure 4)**. The ARGs patterns among the STEC were found to be *blaSHV + tetA* (n = 2)*, CITM + tetA+ aac(3)-IV* (n = 2), *tetA + aac(3)-IV* (n = 2), and *CITM + tetA* (n = 1); that revealed about 43.75% (n = 7/16) STEC isolates carried multiple antibiotic resistant genes **(Figure 4)**.

**Figure 4:**
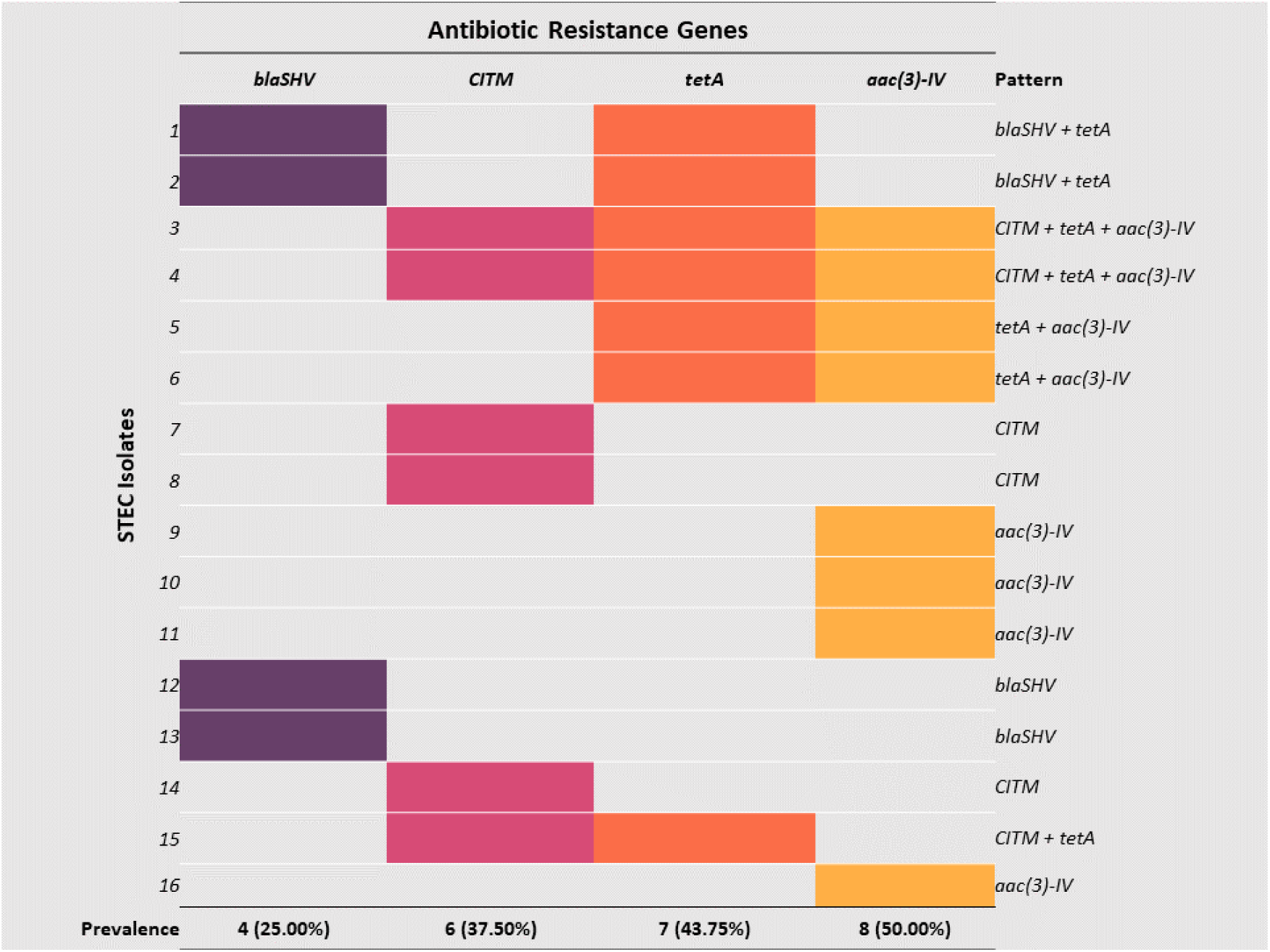
Prevalence of AGRs in STEC samples. (The left side shows 16 different STEC isolates and the right side shows the pattern of AGRs present in each isolate.)

### Antimicrobial susceptibility of STEC

Antimicrobial susceptibility test by disc diffusion method revealed that all STEC isolates (n = 16) were multidrug-resistant **(Figure 5)**. All of them were resistant to amoxicillin (100%). The second highest resistance was recorded against ampicillin (87.50%), followed by gentamycin (75%), tetracycline (68.75%), vancomycin (68.75%), and novobiocin (68.75%) **(Figure 5)**. On the other hand, STEC isolates were mostly sensitive to aztreonam (81.25%) and meropenem (68.75%) **(Figure 5)**.

**Figure 5:**
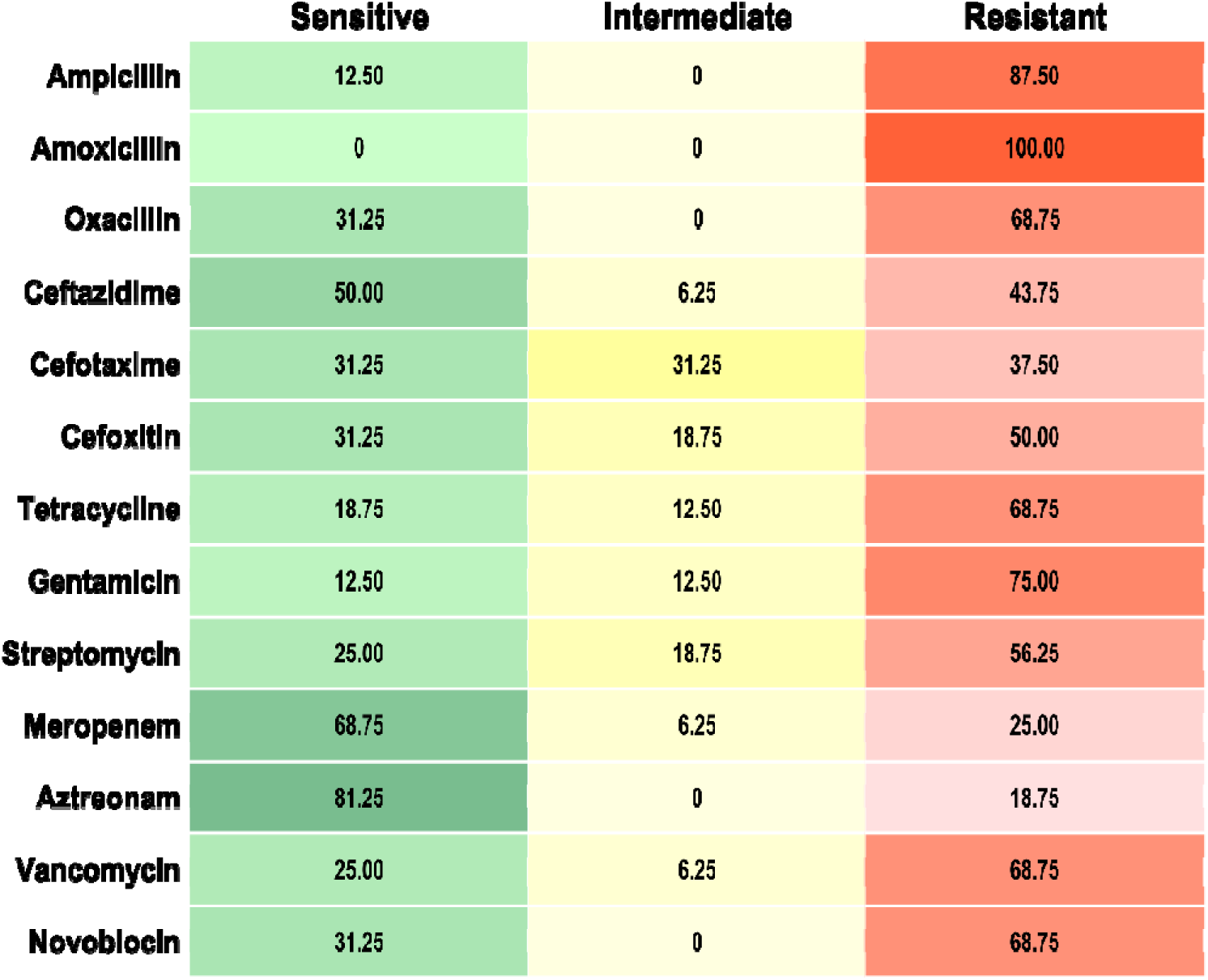
Antibiotic resistance profile of STEC isolates (n = 16) isolated from SCM milk.

### Phylogenetic analysis of MDR-STEC isolate

Phylogenetic analysis of amplified DNA sequence of the *stx1* gene of MDR-STEC revealed that our study sequence (NCBI accession no.: OR088923) was closely related to the *stx1 gene* of various *E. coli* serotypes in Group E including both O157:H7 and non-O157:H7 isolated from various sources **(Figure 6)**. However, most of the strains with close relativeness with our study sequence were non-O157:H7 and only two were O157:H7. Among a total of 18 of these *E. coli* serotypes, mostly were from human stool source (n = 6) and cattle source (n = 5), followed by beef (n = 2), cattle feces (n = 2), cheese (n = 1), and human clinical sample (n = 2) **(Figure 6)**. Other groups such as groups A, B, C, and D were distantly related to our study sequence compared to group E, whereas group F was the most distant among all the groups **(Figure 6)**.

**Figure 6:**
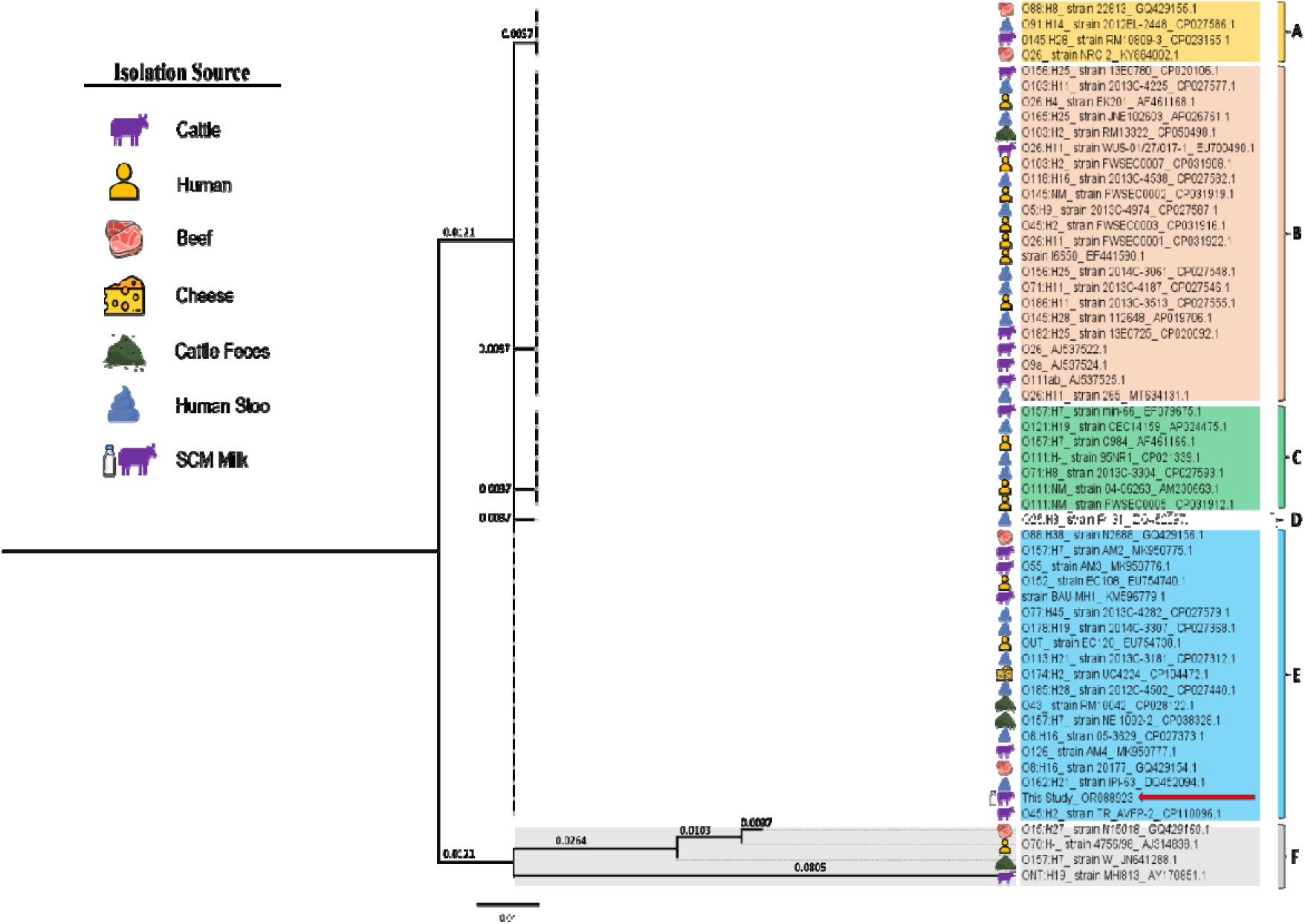
Phylogenetic analysis of *stx1* gene of MDR-STEC isolated from SCM milk. (The red arrow shows the *stx1* gene sequence (NCBI accession no.: OR088923) of the current study. Six different groups (A to F) based on genetic distance are depicted in different colors. The isolation sources of different sequences are illustrated adjacent to the sequence labels.)

## Discussion

Mastitis has a serious financial impact on dairy farming worldwide. SCM is particularly highlighted because there are no obvious visible signs to diagnose (Momtaz et al., 2012). In the current study, we found about 47.2% of the milk samples of crossbreed dairy cattle to be SCM positive with the lowest range of 30% and the highest range of 63.3% **(Figure 2a)**. A similar trend in the prevalence of bovine SCM in Sylhet had been reported in previous studies, where prevalence was found to be 42% to 54% (Das et al., 2020; M. T. Hasan et al., 2017; Kahir et al., 2008; M. M. Rahman et al., 2010). Along with that, SCM is not only highly prevalent in current study areas but also prevalent (51% to 67.9% SCM prevalence) across the country (Md. S. Hasan et al., 2021; Kabir et al., 2017; Kayesh et al., 2014; Rabbani & Samad, 2010). However, in very few regions of Bangladesh, the prevalence of SCM is found to be below 30% (M. A. Islam et al., 2011; Sarker et al., 2013). These previous reports and current study findings indicate that the SCM prevalence remained constant over time, in other words, SCM in dairy cattle is unnoticed or the controlling measures are not being implemented properly.

Moreover, in SCM conditions, the udder appears normal without any detectable clinical signs (Girma & Tamir, 2022). As a result, farmers are unable to isolate the SCM-infected cows and usually, the SCM milk is mixed with normal milk when collected in bulk tanks to supply for human consumption or different milk products manufacturing. Additionally, considering *E. coli* is a predominant environmental bacteria in bovine mastitis condition (Eberhart, 1984; Goulart & Mellata, 2022), and STEC being an emerging bacteria for SCM (Murinda et al., 2019) increases the concern of STEC contamination in consumable milk and dairy products which may pose public health risks. In our study, the number of *E. coli*-positive samples found in cultural and biochemical tests were 110 and 101 out of 184 SCM milk samples, respectively. The cultural and biochemical tests may give false positive results as some non-*E. coli* bacteria can imitate *E. coli* characteristics (Antony et al., 2016). However, molecular detection revealed *E. coli* isolates prevalence of about 50.54% (n = 93 out of 184) in SCM milk **(Figure 2b)**. Previous studies reported that *E. coli* prevalence in SCM milk ranged from 16% to 58.78% (Ahmad et al., 2021; Farhad et al., 2021; Faruk Siddiki, 2019; Haque et al., 2014; Momtaz et al., 2012). Therefore, normal milk mixed with SCM milk which has a predominant presence of *E. coli* is a serious food safety breach that exists in the current dairy food chain in Bangladesh. Furthermore, the concern of contamination gets more complex if there is a presence of STEC.

The STEC is considered to be responsible for causing life-threatening diseases like hemorrhagic colitis and HUS (Smith et al., 2014). STEC is also considered a significant contributor to kidney failure in children (Farrokh et al., 2013). Hence, consumable milk, and milk products that are contaminated with STEC are a serious concern. In our study, 17.20% (n = 16 out of 93) isolates of *E. coli* had at least one STEC virulence gene (*stx1, stx2,* and *eaeA*), and the STEC prevalence in SCM milk was 8.69% (n = 16 out of 184) **(Figure 3a)**. The prevalence of STEC in SCM milk indicates that there is a subtle concern about suffering STEC infection if someone consumes milk or milk products that have been made out of milk that had SCM milk mixed in it. In Bangladesh, 2.05% and 2.15% of STEC prevalence in raw milk were identified (Alam et al., 2021; M. A. Islam et al., 2010). So, SCM milk is more likely than raw milk to be contaminated with STEC. Unfortunately, no reliable study on STEC in SCM milk was found in Bangladesh, but similar studies in other countries reported STEC prevalence in SCM milk as 9% in Egypt (El-Khabaz et al., 2022), 9.7% and 27.23% in Iran (Ahmadi et al., 2020; Momtaz et al., 2012), 9.5% in Brazil (Rangel & Marin, 2009), and 32% in Nigeria (Ivbade et al., 2014). Though our study shows slightly less prevalence of STEC compared to other countries, still this needs to be considered as an emerging threat to public health risks.

Moreover, we detected 31.25% of STEC isolates carrying the *stx1* gene, 56.25% of isolates containing the *stx2* gene, and 6.25% of isolates carrying both *stx1 and stx2* **(Figure 3b).** Additionally, no *eaeA* (intimin) gene was found to be present individually, but 6.25% of isolates carried *stx2 and eaeA* combinedly **(Figure 3b)**. The *stx1 and stx2* genes are the most important virulence factors of STEC, and the *eaeA* gene is considered a complementary virulence factor of STEC in human illness but its mode of action in dairy cattle is yet to be defined (Ahmadi et al., 2020; Mahanti et al., 2013). Among these virulence genes, the *stx2* is more virulent than *stx1*, and the *stx2* is mostly related to hemorrhagic colitis, and HUS (Ahmadi et al., 2020). Our overall study result shows that *stx2* has a higher prevalence in SCM milk **(Figure 3b)**. However, the prevalence of stx2 and stx1 genes has been observed to vary. For instance, a previous study on STEC in SCM milk in Iran reported a higher prevalence of *stx2* (Ahmadi et al., 2020), while another study in the same country found a higher prevalence of *stx1* in STEC isolated from SCM milk (Momtaz et al., 2012). However, in Bangladesh, the *stx2* gene was reported to be predominant in STEC isolated from normal raw milk (Alam et al., 2021). The presence of STEC harboring *stx1,* and *stx2* can cause serious health problems if transmitted to humans. In Bangladesh, enterotoxigenic *E. coli* (ETEC), which is closely linked to STEC (Kaper et al., 2004; Nataro & Kaper, 1998), was estimated to be responsible for 10–20 % of pediatric diarrhea (Qadri et al., 2005). Additionally, outbreaks of STEC that caused HUS in children linked to milk, and milk product consumption have been reported in France, and England (Jones et al., 2019; Treacy et al., 2019). Along with that, if the STEC exhibits antibiotic resistance, then the health and food safety problem may reach a new level of perplexity.

Furthermore, the AMR emergence in animal-origin food and food products due to the use of indiscriminate antibiotics usage is a significant issue for the safety of both human and animal health (Al Amin et al., 2020; Rousham et al., 2018). In the current study, all of the STEC isolates had at least one of the four (*blaSHV, CITM, tet(A), and aac(3)-IV*) ARGs **(Figure 4)**. The most prevalent antibiotic-resistant gene was gentamycin-resistant gene *aac(3)-IV* (50%), followed by tetracycline-resistant *tetA* gene (43.75%), ESBL-resistant *CITM* (37.50%) and *blaSHV* (25%) **(Figure 4)**. Unfortunately, no reliable data could be found on the prevalence of ARGs in STEC isolated from bovine SCM milk in Bangladesh. One study conducted in Bangladesh regarding clinical mastitis milk found the *tet(A)* gene to be present in all the non-STEC *E. coli* isolates but no presence of *blaSHV* (Bag et al., 2021); these findings cannot be compared to our findings as isolates of the previous study were STEC negative (Bag et al., 2021). However, one previous study reported a prevalence of ARGs in STEC isolated from bovine mastitis milk in Iran – *aac(3)-IV* (27.65%), *CITM* (12.76%), and *blaSHV* (6.38%) – which were lower than our findings, but the prevalence of *tet(A)* (48.93%) was higher compared to our study (Momtaz et al., 2012). The ARGs prevalence in STEC may differ due to several factors, such as the predominant use of particular antibiotics in the specific study region; other than that, forage, farm location, management practice, and even frequent contact with other animals can also influence the prevalence of ARGs, as these factors can play a crucial role in adaption and transmission of ARGs (Liu et al., 2024). The spread of antibiotic-resistance genes from *E. coli* pathotypes like STEC to other pathogenic or commensal strains is another aspect of the risk in the public and dairy sectors (Ahmadi et al., 2020; Dormanesh et al., 2014). Moreover, we found STEC isolates were harboring multiple ARGs genes in different combinations such as *blaSHV + tetA, CITM + tetA+ aac(3)-IV*, *tetA + aac(3)-IV,* and *CITM + tetA* **(Figure 4)**. The presence of multiple ARGs in STEC was reported by a previous study as well (Momtaz et al., 2012); the combination pattern or prevalence of different ARGs may change depending on serotypes (Momtaz et al., 2012). In our study, the pattern of ARGs could not be strongly claimed, as only four ARGs were taken into count and no serotyping was performed. However, the presence of multiple ARGs in STEC isolates indicated the presence of multidrug resistance in the identified STEC isolates.

The antimicrobial susceptibility test of STEC isolates in our study showed that all the STEC isolates (n=16 out of 16) were multidrug-resistant (**Figure 5**). We found no studies on the AMR profile of STEC isolated from SCM milk in Bangladesh. A few studies have reported the presence of MDR-*E. coli* in raw milk in Bangladesh (Haque et al., 2014; M. A. Rahman et al., 2017; Rana, Fahim, et al., 2024), and one of these studies reported about 28.13% prevalence (M. A. Rahman et al., 2017). However, outside Bangladesh, a study reported 79.48% MDR-STEC prevalence in bovine SCM milk (Ahmadi et al., 2020). Another study reported 100% multidrug resistance in STEC isolated from bovine mastitis, which supports the current findings (Tavakoli & Pourtaghi, 2017). Hence, we conclude that STEC isolated from SCM milk is predominantly multidrug resistant.

The current study findings also depict that MDR-STEC isolates were fully resistant to amoxicillin and highly resistant to common antibiotics such as ampicillin, gentamycin, and tetracycline (**Figure 5**). A previous study conducted in Bangladesh reported complete resistance of *E. coli* (did not identify STEC) isolated from SCM milk against amoxicillin and ampicillin (Haque et al., 2014). Other studies conducted on *E. coli* isolated from bovine mastitis in Bangladesh, also reported high resistance against tetracycline (89.5%), ampicillin (89.5%), and amoxicillin (94.5%) (Bag et al., 2021). In Iran, a study on STEC from SCM milk reported high resistance to tetracycline and ampicillin, which aligns with the findings of the current study. However, 50% of the STEC were susceptible to gentamicin, though the susceptibility varied depending on the STEC serotypes (Ahmadi et al., 2020). Additionally, a few studies in Bangladesh also reported *E. coli* susceptibility to gentamycin and tetracycline (Farhad et al., 2021; M. A. Islam et al., 2016). On the other hand, a study that took STEC (n = 51) from different sources including human (n = 40), animal (n = 9), and food (n = 2), found that 98.03% (n = 50 out of 51) STEC had resistance against gentamycin (Rubab & Oh, 2020). Based on the current study and previous study findings, it is evident that the commonly used antibiotics have already been resistant, though few previous studies found gentamycin to be susceptible (Ahmadi et al., 2020; Farhad et al., 2021; M. A. Islam et al., 2016). But, in our study, STEC in SCM milk was mostly resistant to gentamycin, which is a concerning issue, because such SCM milk is not regularly checked in dairy farms and is further supplied for human consumption. In this way, resistant pathogens can enter the food chain not only via milk but also through other animal-origin foods, for example, the presence of resistant *E. coli* in beef and poultry has also been reported in Bangladesh (Hossain et al., 2008; Rana, Chowdhury, et al., 2024). Moreover, in our previous studies, we have reported the zoonotic linkage of resistant pathogens in dairy farms (Roy et al., 2024), and also found that biosecurity practices in dairy farms can be potentially associated with human health (Chowdhury et al., 2023). Hence, the possibility of MDR-STEC entering the human food chain via animal source food is obvious if proper safety and precautions are not taken in the first place. These potentially fatal STEC infections may lead to a poor prognosis due to the increase in multidrug resistance in STEC strains (Ahmadi et al., 2020).

Moreover, the current study result depicts that the *stx1* gene of isolated STEC from SCM milk is closely related to different strains of STEC from cattle, humans, food, human stool, and cattle feces origins **(Figure 6)**. These strains were mostly non-O157, except for two O157 strains of cattle and cattle feces origins. Previous studies also reported non-O157 serotypes to be predominant in cattle (Pradel et al., 2000; Torres et al., 2018). Additionally, our study sequence showed close relativeness with a strain BAU-MH1 (KM596779.1), which was isolated from rectal swabs of apparently healthy cattle in Bangladesh (Hassan et al., 2017). We also found similarities with human-origin strains such as O152_EC108 (EU754740.1) and OUT_EC120 (EU754738.1). These strains were isolated from human clinical samples and were associated with diseases (Rui et al., 2010). Moreover, several STEC serotypes found in lactating dairy cattle were reported to be associated with human diseases (Ballem et al., 2020). Additionally, strains such as O88:H38_N2688 (GQ429156.1) and O8:H16_20177 (GQ429154.1), isolated from beef meat, were also found to be closely related to our study sequence (Xia et al., 2010). These findings raise concern for MDR-STEC cross-contamination and zoonoses. The presence of MDR-STEC in milk, with similarities to disease-causing strains, should be considered a potential threat for foodborne pathogen outbreaks if proper precautions are not taken. However, we cannot confirm the zoonotic pathway of MDR-STEC with our current findings. A thorough investigation is needed on the whole dairy supply chain to ensure the pathway of MDR-STEC spreading from milk.

### Conclusions

We found SCM to be highly prevalent in dairy cattle, with a significant presence of STEC. The STEC isolates contained *stx1, stx2,* and *eaeA* virulence genes. All STEC isolates showed multidrug resistance and harbored ARGs as well. Additionally, the isolated MDR-STEC carried a gene that produces the Shiga toxin-1 that was similar to several disease-causing strains linked to cattle, food, and humans. This suggests that SCM milk may serve as a vehicle for the spread of zoonotic MDR-STEC infections. We need to consider this when implementing appropriate control and preventative measures and creating long-term strategies to ensure the safety of bovine milk-derived food.

## Acknowledgments

We would like to express our sincere gratitude to the local dairy farmers for their invaluable cooperation in providing the necessary samples for this study. Our heartfelt thanks also go to the Department of Medicine and the Department of Genetics and Animal Breeding at the Faculty of Veterinary, Animal and Biomedical Sciences, Sylhet Agricultural University, Bangladesh, for generously providing the laboratory facilities and resources that made this research possible.

## Authors’ contribution statement

Conceptualization: TC, MCR, FMAH

Methodology: TC, MCR

Data curation: TC

Formal analysis: TC

Writing original draft: TC

Writing review & editing: FMAH

Project administration: FMAH

Supervision: FMAH

## Ethical review

The study was conducted under the supervision of the Department of Dairy Science, Faculty of Veterinary, Animal and Biomedical Sciences, Sylhet Agricultural University, Sylhet-3100, Bangladesh. International standards considering animal welfare, laboratory hygiene, and ethics were maintained from the start to the end of this study. Approval for conducting the current study was taken from the respective concerned authority of the university.

## Author Approval

All authors have seen and approved the manuscript.

## References

Ahmad, I., Khattak, S., Ali, R., Nawaz, N., Ullah, K., Khan, S. B., Ali, M., Patching, S. G., & Mustafa, M. Z. (2021). Prevalence and molecular characterization of multidrug-resistant Escherichia coli O157: H7 from dairy milk in the Peshawar region of Pakistan. Journal of Food Safety, 41(6), e12941. 10.1111/jfs.12941

Ahmadi, E., Mardani, K., & Amiri, A. (2020). Molecular detection and antimicrobial resistance patterns of Shiga toxigenic Escherichia coli isolated from bovine subclinical mastitis milk samples in Kurdistan, Iran. Archives of Razi Institute, 75(2), 169–177. 10.22092/ARI.2019.124238.1278

Al Amin, M., Hoque, M. N., Siddiki, A. Z., Saha, S., & Kamal, M. M. (2020). Antimicrobial resistance situation in animal health of Bangladesh. Veterinary World, 13(12), 2713– 2727. 10.14202/vetworld.2020.2713-2727

Alam, Md. K.-U., Sarwar, N., Akther, S., Ahmad, M., & Biswas, P. K. (2021). Isolation and molecular characterization of Shigatoxigenic O157 and Non-O157 Escherichia coli in raw milk marketed in Chittagong, Bangladesh. Asian Journal of Dairy and Food Research, 40(1), 1–7. 10.18805/ajdfr.DR-178

Ansharieta, R., Effendi, M. H., & Plumeriastuti, H. (2021). Genetic identification of Shiga toxin encoding gene from cases of multidrug resistance (MDR) Escherichia coli isolated from raw milk. Tropical Animal Science Journal, 44(1), Article 1. 10.5398/tasj.2021.44.1.10

Antony, A., Paul, M., Silvester, R., Aneesa, P. A., Suresh, K., Divya, P. S., Paul, S., Fathima, P. A., & Abdulla, M. (2016). Comparative evaluation of EMB Agar and Hicrome E. coli agar for differentiation of green metallic sheen producing non E. coli and typical E. coli colonies from food and environmental samples. Journal of Pure and Applied Microbiology, 10(4), 2863–2870. 10.22207/JPAM.10.4.48

Astermark, S., Funke, H., & Engan-Skei, I. (1969). The relationship between the California Mastitis Test, Whiteside Test, Brabant Mastitis Reaction, Catalase Test, and Direct Cell Counting of Milk. Acta Veterinaria Scandinavica, 10(2), 146–167. 10.1186/BF03548286

Bag, Md. A. S., Khan, Md. S. R., Sami, Md. D. H., Begum, F., Islam, Md. S., Rahman, Md. M., Rahman, Md. T., & Hassan, J. (2021). Virulence determinants and antimicrobial resistance of E. coli isolated from bovine clinical mastitis in some selected dairy farms of Bangladesh. Saudi Journal of Biological Sciences, 28(11), 6317–6323. 10.1016/j.sjbs.2021.06.099

Ballem, A., Gonçalves, S., Garcia-Meniño, I., Flament-Simon, S. C., Blanco, J. E., Fernandes, C., Saavedra, M. J., Pinto, C., Oliveira, H., Blanco, J., Almeida, G., & Almeida, C. (2020). Prevalence and serotypes of Shiga toxin-producing Escherichia coli (STEC) in dairy cattle from Northern Portugal. PLOS ONE, 15(12), e0244713. 10.1371/journal.pone.0244713

Bauer, A. W., Kirby, W. M. M., Sherris, J. C., & Turck, M. (1966). Antibiotic susceptibility testing by a standardized single disk method. American Journal of Clinical Pathology, 45(4_ts), 493–496. 10.1093/ajcp/45.4_ts.493

Beveridge, T. (2001). Use of the Gram stain in microbiology. Biotechnic & Histochemistry, 76(3), 111–118. 10.1080/bih.76.3.111.118

Chowdhury, T., Ahmed, J., Hossain, M. T., Roy, M. C., Ashik-Uz-Zaman, M., Uddin, M. N., Rahman, M. M., Kabir, M. G., & Hossain, F. M. A. (2023). Knowledge, attitudes and biosecurity practices among the small-scale dairy farmers in Sylhet district, Bangladesh. Veterinary Medicine and Science, 1-9. 10.1002/vms3.1199

CLSI. (2020). Performance Standards for Antimicrobial Susceptibility Testing (30th ed.). CLSI supplement M100. Wayne, PA: Clinical and Laboratory Standards Institute;

Daini, O. A., Ogbolu, O. D., & Ogunledun, A. (2005). Quinolones resistance and R-plasmids of some gram negative enteric bacilli. African Journal of Clinical and Experimental Microbiology, 6(1), Article 1. 10.4314/ajcem.v6i1.7394

Das, M., Barman, A., Akter, S., Chowdhury, S. R., Ashik-Uz-Zaman, M., & Rahman, M. M. (2020). Prevalence of subclinical mastitis and physicochemical properties of milk sample in dairy cows of Sylhet district. International Journal of Natural Sciences, 10(2), 30–36.

Dash, S. K., Chakraborty, S. P., & Roy, S. (2012). Isolation and characterization of multi drug resistant uropathogenic Escherichia coli from urine sample of urinary tract infected patients. 2(1), 15.

Dipineto, L., Santaniello, A., Fontanella, M., Lagos, K., Fioretti, A., & Menna, L. f. (2006). Presence of Shiga toxin-producing Escherichia coli O157:H7 in living layer hens. Letters in Applied Microbiology, 43(3), 293–295. 10.1111/j.1472-765X.2006.01954.x

Dormanesh, B., Safarpoor Dehkordi, F., Hosseini, S., Momtaz, H., Mirnejad, R., Hoseini, M. J., Yahaghi, E., Tarhriz, V., & Khodaverdi Darian, E. (2014). Virulence factors and o-serogroups profiles of uropathogenic Escherichia coli isolated from Iranian pediatric patients. Iranian Red Crescent Medical Journal, 16(2), e14627. 10.5812/ircmj.14627

Eberhart, R. J. (1984). Coliform mastitis. The Veterinary Clinics of North America. Large Animal Practice, 6(2), 287–300. 10.1016/s0196-9846(17)30023-x

Elafify, M., Khalifa, H. O., Al-Ashmawy, M., Elsherbini, M., El Latif, A. A., Okanda, T., Matsumoto, T., Koseki, S., & Abdelkhalek, A. (2020). Prevalence and antimicrobial resistance of Shiga toxin-producing Escherichia coli in milk and dairy products in Egypt. Journal of Environmental Science and Health, Part B, 55(3), 265–272. 10.1080/03601234.2019.1686312

Elhadidy, M., & Mohammed, M. A. (2013). Shiga toxin-producing *Escherichia coli* from raw milk cheese in Egypt: Prevalence, molecular characterization and survival to stress conditions. Letters in Applied Microbiology, 56(2), 120–127. 10.1111/lam.12023

El-Khabaz, K. A. S., Elshrief, L. M. T., & Elmeligy, E. (2022). Genetic assessment of Shiga toxin and antibiotic resistance of E. coli isolated from milk of cows infected with sub-clinical mastitis. Journal of Advanced Veterinary Research, 12(3), Article 3.

Fahim, K. M., Ismael, E., Khalefa, H. S., Farag, H. S., & Hamza, D. A. (2019). Isolation and characterization of *E. coli* strains causing intramammary infections from dairy animals and wild birds. International Journal of Veterinary Science and Medicine, 7(1), 61–70. 10.1080/23144599.2019.1691378

Farhad, S., Shahid, M., Mahmud, M., Kabir, A., Das, S., Rahman, M., & Nazir, K. (2021). Molecular detection and antibiogram of Escherichia coli O157 isolated from subclinical mastitis affected cows at Baghabari, Sirajganj. Veterinary Research Notes, 1(2), 6. 10.5455/vrn.2021.a2

Farrokh, C., Jordan, K., Auvray, F., Glass, K., Oppegaard, H., Raynaud, S., Thevenot, D., Condron, R., De Reu, K., Govaris, A., Heggum, K., Heyndrickx, M., Hummerjohann, J., Lindsay, D., Miszczycha, S., Moussiegt, S., Verstraete, K., & Cerf, O. (2013). Review of Shiga-toxin-producing Escherichia coli (STEC) and their significance in dairy production. International Journal of Food Microbiology, 162(2), 190–212. 10.1016/j.ijfoodmicro.2012.08.008

Faruk Siddiki, S. H. M. (2019). Comparison of bacterial pathogens associated with different types of bovine mastitis and their antibiotic resistancestatus in bangladesh. Journal of Veterinary Medical and One Health Research, 1(1), 17–27. 10.36111/jvmohr.2019.1(1).0002

Feng, P., Weagant, S. D., & Jinneman, K. (2020). BAM Chapter 4A: Diarrheagenic Escherichia coli. In Bacteriological analytical manual (BAM). FDA. https://www.fda.gov/food/laboratory-methods-food/bam-chapter-4a-diarrheagenic-escherichia-coli

Girma, A., & Tamir, D. (2022). Prevalence of bovine mastitis and its associated risk factors among dairy cows in ethiopia during 2005–2022: A systematic review and meta-analysis. Veterinary Medicine International, 2022(Article ID 7775197), 1–19. 10.1155/2022/7775197

Gorman, R., & Adley, C. C. (2004). An evaluation of five preservation techniques and conventional freezing temperatures of −20oC and −85oC for long-term preservation of Campylobacter jejuni. Letters in Applied Microbiology, 38(4), 306–310. 10.1111/j.1472-765X.2004.01490.x

Goulart, D. B., & Mellata, M. (2022). Escherichia coli mastitis in dairy cattle: etiology, diagnosis, and treatment challenges. Frontiers in Microbiology, 13(Article 928346). https://www.frontiersin.org/articles/10.3389/fmicb.2022.928346

Guillaume, G., Verbrugge, D., Chasseur-Libotte, M.-L., Moens, W., & Collard, J.-M. (2000). PCR typing of tetracycline resistance determinants (Tet A–E) in Salmonella enterica serotype Hadar and in the microbial community of activated sludges from hospital and urban wastewater treatment facilities in Belgium. FEMS Microbiology Ecology, 32(1), 77–85. 10.1111/j.1574-6941.2000.tb00701.x

Gyles, C. L. (2007). Shiga toxin-producing Escherichia coli: An overview1. Journal of Animal Science, 85(suppl_13), E45–E62. 10.2527/jas.2006-508

Haque, M. E., Islam, M. A., Akter, S., & Saha, S. (2014). Identification, molecular detection and antibiogram profile of bacteria isolated from California Mastitis test positive milk samples of crossbred cows of Satkhira District in Bangladesh. GSTF Journal of Veterinary Science (JVet), 1(1), Article 1.

Hasan, M. T., Islam, M. R., Runa, N. S., Hasan, M., N., Uddin, A. H. M. M., & Singh, S. K. (2017). Study on bovine sub-clinical mastitis on farm condition with special emphasis on antibiogram of the causative bacteria. Bangladesh Journal of Veterinary Medicine, 14(2), 161–166. 10.3329/bjvm.v14i2.31386

Hasan, Md. S., Humayun Kober, A. K. M., Ahmed Rana, E., & Saiful Bari, M. (2021). Association of udder lesions with subclinical mastitis in dairy cows of chattogram, bangladesh. Advances in Animal and Veterinary Sciences, 10(2), 226–235. 10.17582/journal.aavs/2022/10.2.226.235

Hassan, J., Nazir, K., Parvej, M., Kamal, T., & Rahman, M. (2017). Molecular based prevalence of shigatoxigenic Escherichia coli in rectal swab of apparently healthy cattle in Mymensingh district, Bangladesh. Journal of Advanced Veterinary and Animal Research, 4(2), 1. 10.5455/javar.2017.d213

Hessain, A. M., Al-Arfaj, A. A., Zakri, A. M., El-Jakee, J. K., Al-Zogibi, O. G., Hemeg, H. A., & Ibrahim, I. M. (2015). Molecular characterization of *Escherichia coli* O157:H7 recovered from meat and meat products relevant to human health in Riyadh, Saudi Arabia. Saudi Journal of Biological Sciences, 22(6), 725–729. 10.1016/j.sjbs.2015.06.009

Hinthong, W., Pumipuntu, N., Santajit, S., Kulpeanprasit, S., Buranasinsup, S., Sookrung, N., Chaicumpa, W., Aiumurai, P., & Indrawattana, N. (2017). Detection and drug resistance profile of Escherichia coli from subclinical mastitis cows and water supply in dairy farms in Saraburi Province, Thailand. PeerJ, 5, e3431. 10.7717/peerj.3431

Hossain, F. Mohd. A., Hossain, Md. M., & Hossain, Md. T. (2011). Antibiogram profile of Escherichia coli isolated from migratory birds. Eurasian Journal of Veterinary Sciences, 27(3), 167–170.

Hossain, M., Siddique, M., Hossain, F., Zinnah, M., Hossain, M., Alam, M., Rahman, M., & Choudhury, K. (2008). Isolation, identification, toxin profile and antibiogram of *Escherichia coli* isolated from broilers and layers in Mymensingh district of Bangladesh. Bangladesh Journal of Veterinary Medicine, 6(1), 1–5. 10.3329

Islam, M. A., Islam, M. Z., Islam, M. A., Rahman, M. S., & Islam, M. T. (2011). Prevalence of subclinical mastitis in dairy cows in selected areas of Bangladesh. Bangladesh Journal of Veterinary Medicine, 9(1), Article 1. 10.3329/bjvm.v9i1.11216

Islam, M. A., Kabir, S. M. L., & Seel, S. K. (2016). Molecular detection and characterization of escherichia coli isolated from raw milk sold in different markets of bangladesh. Bangladesh Journal of Veterinary Medicine, 14(2), Article 2. 10.3329/bjvm.v14i2.31408

Islam, M. A., Mondol, A. S., Azmi, I. J., de Boer, E., Beumer, R. R., Zwietering, M. H., Heuvelink, A. E., & Talukder, K. A. (2010). Occurrence and characterization of Shiga Toxin–Producing Escherichia coli in raw meat, raw milk, and street vended juices in Bangladesh. Foodborne Pathogens and Disease, 7(11), 1381–1385. 10.1089/fpd.2010.0569

Islam, Md. S., Nayeem, Md. M. H., Sobur, Md. A., Ievy, S., Islam, Md. A., Rahman, S., Kafi, Md. A., Ashour, H. M., & Rahman, Md. T. (2021). Virulence determinants and multidrug resistance of Escherichia coli isolated from migratory birds. Antibiotics, 10(2), 190. 10.3390/antibiotics10020190

Ivbade, A., Ojo, O. E., & Dipeolu, M. A. (2014). Shiga toxin-producing Escherichia coli O157:H7 in milk and milk products in Ogun State, Nigeria. Veterinaria Italiana, 50(3), 185–191. 10.12834/VetIt.129.2187.1

Jones, G., Lefèvre, S., Donguy, M.-P., Nisavanh, A., Terpant, G., Fougère, E., Vaissière, E., Guinard, A., Mailles, A., de Valk, H., Fila, M., Tanné, C., Le Borgne, C., Weill, F.-X., Bonacorsi, S., Jourdan-Da Silva, N., & Mariani-Kurkdjian, P. (2019). Outbreak of Shiga toxin-producing Escherichia coli (STEC) O26 paediatric haemolytic uraemic syndrome (HUS) cases associated with the consumption of soft raw cow’s milk cheeses, France, March to May 2019. Euro Surveillance: Bulletin Europeen Sur Les Maladies Transmissibles = European Communicable Disease Bulletin, 24(22). 10.2807/1560-7917.ES.2019.24.22.1900305

Kabir, M. H., Ershaduzzaman, M., Giasuddin, M., Islam, M. R., Nazir, K. H. M. N. H., Islam, M. S., Karim, M. R., Rahman, M. H., & Ali, M. Y. (2017). Prevalence and identification of subclinical mastitis in cows at BLRI Regional Station, Sirajganj, Bangladesh. Journal of Advanced Veterinary and Animal Research, 4(3), Article 3.

Kahir, M. A., Islam, M. M., Rahman, A. K. M. A., Nahar, A., Rahman, M. S., & Son, H.-J. (2008). Prevalence and risk factors of subclinical bovine mastitis in some dairy farms of Sylhet district of Bangladesh. Korean Journal of Veterinary Service, 31(4), 497– 504.

Kalyaanamoorthy, S., Minh, B. Q., Wong, T. K. F., von Haeseler, A., & Jermiin, L. S. (2017). ModelFinder: Fast model selection for accurate phylogenetic estimates. Nature Methods, 14(6), 587–589. 10.1038/nmeth.4285

Kaper, J. B., Nataro, J. P., & Mobley, H. L. T. (2004). Pathogenic Escherichia coli. Nature Reviews Microbiology, 2(2), Article 2. 10.1038/nrmicro818

Karas, J. A., Pillay, D. G., & Sturm, A. W. (2007). The catalase reaction of *Shigella* species and its use in rapid screening for epidemic *Shigella dysenteriae* type 1. Annals of Tropical Medicine & Parasitology, 101(1), 79–84. 10.1179/136485907154575

Karim, S. J. I., Islam, M., Sikder, T., Rubaya, R., Halder, J., & Alam, J. (2020). Multidrug-resistant Escherichia coli and Salmonella spp. Isolated from pigeons. Veterinary World, 13(10), 2156–2165. 10.14202/vetworld.2020.2156-2165

Kayesh, M., Talukder, M., & Anower, A. (2014). Prevalence of subclinical mastitis and its association with bacteria and risk factors in lactating cows of Barisal district in Bangladesh. International Journal of Biological Research, 2(2), 35–38. 10.14419/ijbr.v2i2.2835

Khan, A., Yamasaki, S., Sato, T., Ramamurthy, T., Pal, A., Datta, S., Chowdhury, N. R., Das, S. C., Sikdar, A., Tsukamoto, T., Bhattacharya, S. K., Takeda, Y., & Nair, G. B. (2002). Prevalence and genetic profiling of virulence determinants of non-o157 Shiga toxin-producing Escherichia coli isolated from cattle, beef, and humans, Calcutta, India. Emerging Infectious Diseases Journal, 8(1). 10.3201/eid0801.010104

Liu, T., Lee, S., Kim, M., Fan, P., Boughton, R. K., Boucher, C., & Jeong, K. C. (2024). A study at the wildlife-livestock interface unveils the potential of feral swine as a reservoir for extended-spectrum β-lactamase-producing *Escherichia coli*. Journal of Hazardous Materials, 473, 134694. 10.1016/j.jhazmat.2024.134694

Magiorakos, A.-P., Srinivasan, A., Carey, R. B., Carmeli, Y., Falagas, M. E., Giske, C. G., Harbarth, S., Hindler, J. F., Kahlmeter, G., Olsson-Liljequist, B., Paterson, D. L., Rice, L. B., Stelling, J., Struelens, M. J., Vatopoulos, A., Weber, J. T., & Monnet, D. L. (2012). Multidrug-resistant, extensively drug-resistant and pandrug-resistant bacteria: An international expert proposal for interim standard definitions for acquired resistance. Clinical Microbiology and Infection, 18(3), 268–281. 10.1111/j.1469-0691.2011.03570.x

Mahanti, A., Samanta, I., Bandopaddhay, S., Joardar, S. N., Dutta, T. K., Batabyal, S., Sar, T. K., & Isore, D. P. (2013). Isolation, molecular characterization and antibiotic resistance of Shiga Toxin-Producing Escherichia coli (STEC) from buffalo in India. Letters in Applied Microbiology, 56(4), 291–298. 10.1111/lam.12048

Momtaz, H., Safarpoor Dehkordi, F., Taktaz, T., Rezvani, A., & Yarali, S. (2012). Shiga Toxin-Producing Escherichia coli isolated from bovine mastitic milk: serogroups, virulence factors, and antibiotic resistance properties. The Scientific World Journal, 2012(Article 618709), 1–9. 10.1100/2012/618709

Murinda, S. E., Ibekwe, A. M., Rodriguez, N. G., Quiroz, K. L., Mujica, A. P., & Osmon, K. (2019). Shiga Toxin–Producing Escherichia coli in mastitis: An international perspective. Foodborne Pathogens and Disease, 16(4), 229–243. 10.1089/fpd.2018.2491

Mylius, M., Dreesman, J., Pulz, M., Pallasch, G., Beyrer, K., Claußen, K., Allerberger, F., Fruth, A., Lang, C., Prager, R., Flieger, A., Schlager, S., Kalhöfer, D., & Mertens, E. (2018). Shiga toxin-producing Escherichia coli O103:H2 outbreak in Germany after school trip to Austria due to raw cow milk, 2017 – The important role of international collaboration for outbreak investigations. International Journal of Medical Microbiology, 308(5), 539–544. 10.1016/j.ijmm.2018.05.005

Nataro, J. P., & Kaper, J. B. (1998). Diarrheagenic Escherichia coli. Clinical Microbiology Reviews, 11(1), 142–201. 10.1128/CMR.11.1.142

Nguyen, L.-T., Schmidt, H. A., von Haeseler, A., & Minh, B. Q. (2015). IQ-TREE: A fast and effective stochastic algorithm for estimating maximum-likelihood phylogenies. Molecular Biology and Evolution, 32(1), 268–274. 10.1093/molbev/msu300

Nourbakhsh, F., Rajai, M., & Momtaz, H. (2017). Antibiotic resistance and carriage integron classes in clinical isolates of Acinetobacter Baumannii from Isfahan Hospitals, Iran. Zahedan Journal of Research in Medical Sciences, 19(1), Article 1. 10.17795/zjrms-6009

Parvin, M. S., Talukder, S., Ali, M. Y., Chowdhury, E. H., Rahman, M. T., & Islam, M. T. (2020). Antimicrobial resistance pattern of Escherichia coli isolated from frozen chicken meat in Bangladesh. Pathogens, 9(6), Article 6. 10.3390/pathogens9060420

Pradel, N., Livrelli, V., De Champs, C., Palcoux, J.-B., Reynaud, A., Scheutz, F., Sirot, J., Joly, B., & Forestier, C. (2000). Prevalence and characterization of Shiga Toxin-Producing Escherichia coli isolated from cattle, food, and children during a one-year prospective study in France. Journal of Clinical Microbiology, 38(3), 1023–1031.

Purkayastha, M., Khan, M., Alam, M., Siddique, M., Begum, F., Mondal, T., & Choudhury, S. (1970). Cultural and biochemical characterization of sheep Escherichia coli isolated from in and around BAU campus. Bangladesh Journal of Veterinary Medicine, 8(1), 51–55. 10.3329/bjvm.v8i1.8350

Qadri, F., Svennerholm, A.-M., Faruque, A. S. G., & Sack, R. B. (2005). Enterotoxigenic Escherichia coli in developing countries: Epidemiology, microbiology, clinical features, treatment, and prevention. Clinical Microbiology Reviews, 18(3), 465–483. 10.1128/CMR.18.3.465-483.2005

Rana, S., Chowdhury, K., Muchemi, J., Fahim, F. J., Das, R., Ali, M., Noor, M., Zinnah, K. M. A., Hussain, S. N., & Hossain, F. M. A. (2024). Molecular characterization of biofilm producing *Escherichia coli* isolated from beef value chain in Bangladesh. Food Science of Animal Products, 2(2), 9240059–9240059. 10.26599/FSAP.2024.9240059

Rana, S., Fahim, F. J., Das, R., Abdullah, K. S., Islam, S. S., Mahim, N. J., Sultana, N., Uddin, M. N., Islam, M. R., Ahmad, S., Zinnah, K., Noor, M., & Hossain, F. M. A. (2024). High prevalence of multidrug-resistant extended spectrum beta lactamase-producing Escherichia coli in raw milk in Bangladesh. Microbes and Infectious Diseases. 10.21608/mid.2024.258318.1735

Rabbani, A., & Samad, M. A. (2010). Host determinants based comparative prevalence of subclinical mastitis in lactating holstein-friesian cross cows and red chittagong cows in bangladesh. Bangladesh Journal of Veterinary Medicine, 8(1), Article 1. 10.3329/bjvm.v8i1.7397

Rahman, M. A., Rahman, A. K. M. A., Islam, M. A., & Alam, M. M. (2017). Antimicrobial resistance of escherichia coli isolated from milk, beef and chicken meat in bangladesh. Bangladesh Journal of Veterinary Medicine, 15(2), Article 2. 10.3329/bjvm.v15i2.35525

Rahman, M. M., Islam, M. R., Uddin, M. B., & Aktaruzzaman, M. (2010). Prevalence of subclinical mastitis in dairy cows reared in sylhet District of Bangladesh. International Journal of Biological Research, 1(2), 23–28.

Rangel, P., & Marin, J. M. (2009). Analysis of Escherichia coli isolated from bovine mastitic milk. Pesquisa Veterinária Brasileira, 29(5), 363–368. 10.1590/S0100-736X2009000500001

Reinders, R. d., Barna, A., Lipman, L. j. a., & Bijker, P. g. h. (2002). Comparison of the sensitivity of manual and automated immunomagnetic separation methods for detection of Shiga toxin-producing Escherichia coli O157:H7 in milk. Journal of Applied Microbiology, 92(6), 1015–1020. 10.1046/j.13652672.2002.01646.x

Rousham, E. K., Unicomb, L., & Islam, M. A. (2018). Human, animal and environmental contributors to antibiotic resistance in low-resource settings: Integrating behavioural, epidemiological and One Health approaches. Proceedings. Biological Sciences, 285(1876), 20180332. 10.1098/rspb.2018.0332

Rozwadowski, M., & Gawel, D. (2022). Molecular factors and mechanisms driving multidrug resistance in uropathogenic Escherichia coli—An Update. Genes, 13(8), 1397. 10.3390/genes13081397

Roy, M. C., Chowdhury, T., Hossain, M. T., Hasan, Md. M., Zahran, E., Rahman, Md. M., Zinnah, K. M. A., Rahman, Md. M., & Hossain, F. M. A. (2024). Zoonotic linkage and environmental contamination of Methicillin-resistant *Staphylococcus aureus* (MRSA) in dairy farms: A one health perspective. One Health, 18, 100680. 10.1016/j.onehlt.2024.100680

Rubab, M., & Oh, D.-H. (2020). Virulence characteristics and antibiotic resistance profiles of Shiga Toxin-Producing Escherichia coli isolates from diverse sources. Antibiotics, 9(9), 587. 10.3390/antibiotics9090587

Rui, L., Takaaki, H., Ken-ichi, H., & Takashia, M. (2010). Phylogenetic analysis and Shiga toxin production profiling of Shiga toxin-producing/enterohemorrhagic Escherichia coli clinical isolates. Microbial Pathogenesis, 49(5). 10.1016/j.micpath.2010.06.005

Sanger, F., Nicklen, S., & Coulson, A. R. (1977). DNA sequencing with chain-terminating inhibitors. Proceedings of the National Academy of Sciences of the United States of America, 74(12), 5463–5467.

Sarker, S. C., Parvin, Mst. S., Rahman, A. K. M. A., & Islam, Md. T. (2013). Prevalence and risk factors of subclinical mastitis in lactating dairy cows in north and south regions of Bangladesh. Tropical Animal Health and Production, 45(5), 1171–1176. 10.1007/s11250-012-0342-7

Smith, J. L., Fratamico, P. M., & Gunther, N. W. (2014). Chapter Three—Shiga toxin-producing Escherichia coli. In S. Sariaslani & G. M. Gadd (Eds.), Advances in Applied Microbiology (Vol. 86, pp. 145–197). Academic Press. 10.1016/B978-0-12-800262-9.00003-2

Sweeney, M. T., Lubbers, B. V., Schwarz, S., & Watts, J. L. (2018). Applying definitions for multidrug resistance, extensive drug resistance and pandrug resistance to clinically significant livestock and companion animal bacterial pathogens. Journal of Antimicrobial Chemotherapy, 73(6), 1460–1463. 10.1093/jac/dky043

Tarr, P. I., Gordon, C. A., & Chandler, W. L. (2005). Shiga-toxin-producing Escherichia coli and haemolytic uraemic syndrome. Lancet (London, England), 365(9464), 1073– 1086. 10.1016/S0140-6736(05)71144-2

Tavakoli, M., & Pourtaghi, H. (2017). Molecular detection of virulence genes and multi-drug resistance patterns in Escherichia coli (STEC) in clinical bovine mastitis: Alborz province, Iran. Iranian Journal of Veterinary Research, 18(3), 208–211.

Torres, A. G., Amaral, M. M., Bentancor, L., Galli, L., Goldstein, J., Krüger, A., & Rojas-Lopez, M. (2018). Recent advances in Shiga toxin-producing Escherichia coli research in Latin America. Microorganisms, 6(4), Article 4. 10.3390/microorganisms6040100

Treacy, J., Jenkins, C., Paranthaman, K., Jorgensen, F., Mueller-Doblies, D., Anjum, M., Kaindama, L., Hartman, H., Kirchner, M., Carson, T., & Kar-Purkayastha, I. (2019). Outbreak of Shiga toxin-producing Escherichia coli O157:H7 linked to raw drinking milk resolved by rapid application of advanced pathogen characterisation methods, England, August to October 2017. Eurosurveillance, 24(16), 1800191. 10.2807/1560-7917.ES.2019.24.16.1800191

Wallace, J. S., Cheasty, T., & Jones, K. (1997). Isolation of Vero cytotoxin-producing Escherichia coli O157 from wild birds. Journal of Applied Microbiology, 82(3), 399–404. 10.1046/j.1365-2672.1997.00378.x

Xia, X., Meng, J., McDermott, P. F., Ayers, S., Blickenstaff, K., Tran, T.-T., Abbott, J., Zheng, J., & Zhao, S. (2010). Presence and characterization of shiga toxin-producing Escherichia coli and other potentially diarrheagenic E. coli strains in retail meats. Applied and Environmental Microbiology, 76(6), 1709–1717. 10.1128/AEM.01968-09

