## Supplementary Table 1 for "Prevalence and zoonotic risk of multidrug-resistant Escherichia coli in bovine subclinical mastitis milk: Insights into the virulence and antimicrobial resistance"

**Supplementary Table 1: List of antibiotics used for antimicrobial susceptibility test.**

| Antibiotic Groups | Names of Antibiotics |
| --- | --- |
| Penicillin | Ampicillin (AMP, 10μg), Amoxicillin (AMX, 30μg), Oxacillin (OX, 1μg) |
| Cephalosporins | Ceftazidime (CAZ, 30μg), Cefotaxime (CTX, 30μg), Cefoxitin (CX, 30μg) |
| Tetracyclines | Tetracycline (TE, 30μg) |
| Aminoglycosides | Gentamicin (GEN, 10μg), Streptomycin (S, 10μg) |
| Carbapenem | Meropenem (MEM, 10μg) |
| Monobactam | Aztreonam (AT, 30μg) |
| Glycopeptide | Vancomycin (VA, 30μg) |
| Aminocoumarin | Novobiocin (NV, 30µg) |
