## Supplementary Table 2 for "Prevalence and zoonotic risk of multidrug-resistant Escherichia coli in bovine subclinical mastitis milk: Insights into the virulence and antimicrobial resistance"

**Supplementary Table 2: Different characteristics of presumptively confirmed *E. coli* isolates found in cultural media and biochemical tests in the current study.**

| **Cultural and Biochemical tests** | | **Characteristics of *E. coli* isolates** |
| --- | --- | --- |
| EMB | | Colonies with dark center and green metallic sheen |
| Gram Staining | | Gram (-ve), rod-shaped, pink color organism |
| Motility Indole Urea test | Motility test | +ve (uniform turbidity in medium) |
|  | Indole test | +ve (cherry-red color ring) |
|  | Urea test | -ve |
| Methyl Red test | | +ve (pink-red color development) |
| Voges-Proskauer test | | -ve (no color change) |
| Citrate test | | -ve (no change of color) |
| Growth on Sugar fermentation test (TSI) | Butt | Yellow/Acidic |
|  | Slant | Yellow/Acidic |
|  | Gas | +ve (presence of gas bubble) |
|  | H_2_S | -ve |
| Catalase test | | +ve |
